# MetaEuk – sensitive, high-throughput gene discovery and annotation for large-scale eukaryotic metagenomics

**DOI:** 10.1101/851964

**Authors:** Eli Levy Karin, Milot Mirdita, Johannes Söding

## Abstract

**Background:** Metagenomics is revolutionizing the study of microorganisms and their involvement in biological, biomedical, and geochemical processes, allowing us to investigate by direct sequencing a tremendous diversity of organisms without the need for prior cultivation. Unicellular eukaryotes play essential roles in most microbial communities as chief predators, decomposers, phototrophs, bacterial hosts, symbionts and parasites to plants and animals. Investigating their roles is therefore of great interest to ecology, biotechnology, human health, and evolution. However, the generally lower sequencing coverage, their more complex gene and genome architectures, and a lack of eukaryote-specific experimental and computational procedures have kept them on the sidelines of metagenomics.

**Results:** MetaEuk is a toolkit for high-throughput, reference-based discovery and annotation of protein-coding genes in eukaryotic metagenomic contigs. It performs fast searches with 6-frame-translated fragments covering all possible exons and optimally combines matches into multi-exon proteins. We used a benchmark of seven diverse, annotated genomes to show that MetaEuk is highly sensitive even under conditions of low sequence similarity to the reference database. To demonstrate MetaEuk’s power to discover novel eukaryotic proteins in large-scale metagenomic data, we assembled contigs from 912 samples of the Tara Oceans project. MetaEuk predicted >12,000,000 protein-coding genes in eight days on ten 16-core servers. Most of the discovered proteins are highly diverged from known proteins and originate from very sparsely sampled eukaryotic supergroups.

**Conclusion:** The open-source (GPLv3) MetaEuk software (https://github.com/soedinglab/metaeuk) enables large-scale eukaryotic metagenomics through reference-based, sensitive taxonomic and functional annotation.

## Background

Unicellular eukaryotes are present in almost all environments, including soil [1], oceans [2], and plant and animal-associated microbiomes [3, 4]. They exhibit both autotrophic and heterotrophic lifestyles [5], exist in symbiosis with plants and animals [6], and interact with other microbial organisms [7]. They account for roughly half of the global primary productivity in the oceans, mostly by photosynthesis [8], are key contributors to the carbon and nitrogen cycles through carbon-dioxide fixation, organic matter degradation, and denitrification [9, 10], and have been shown to be a source for chemically bioactive compounds [e.g., 11,12].

Since the advent of metabarcoding using 18S rRNA genes, the known evolutionary diversity of unicellular eukaryotes has increased by orders of magnitude [13], and novel phyla and supra-kingdoms are still being discovered [14, 15]. Due to their vast diversity [16, 17], unicellular eukaryotes are certain to hold invaluable secrets for biotechnology and biomedicine.

Protein-coding genes are major keys for understanding eukaryotic functions and activities [18]. Metatranscriptomic and metagenomic studies provide unique means to reveal protein-coding genes. However, despite the great potential of studying uncultivatable eukaryotes in their natural environment, they have received little attention in metatranscriptomic and metagenomic studies so far, with a few notable exceptions [e.g., 19,20]. The unique features of eukaryotic data, i.e., lower genomic coverage due to lower population densities in metagenomic samples, fewer reference genomes, increased genome sizes and higher complexity of gene structure negatively impact all stages of metagenomic analyses, from assembly, through binning, to protein prediction and annotation [as discussed by 21,22].

Specifically, identifying protein-coding genes in eukaryotes is inherently more challenging than in prokaryotes due to the exon-intron architecture of eukaryotic genes. To date, methods for eukaryotic gene calling [e.g., 23–25] consider two types of information when training models for gene prediction: intrinsic sequence signals (e.g., CpG islands) and extrinsic data, such as transcriptomics or an annotated genome from a closely-related organism. As splicing signatures are not well conserved throughout evolution, the predictive power of the trained models declines fast when applied to organisms that are phylogenetically distant from the organism on which the model was trained [26].

While these methods are very useful for genomics, their applicability to metagenomic data is severely limited. First, the transcriptomic or genomic data of annotated organisms that are sufficiently closely related are usually not available. Second, since the models need to be trained on a relatively narrow clade, the application of such methods to metagenomic data requires to first bin the assembled contigs by their assumed genome of origin [as performed by 27], which is often quite inaccurate and slow, especially when the number of contigs is large, the coverage is low, the contigs are short, and the metagenomic data are species-rich [28–30]. Finally, model-training in itself is time consuming, taking hours to days per genomic bin [25, 27], limiting this approach to the analysis of few genomic bins at a time.

Previously, methods that bypass or reduce the need to explicitly train models to detect protein-coding genes have been proposed in the context of genomics [e.g., 31,32]. These methods extract putative protein-coding fragments from the genome and join those that bear sequence similarity to available transcriptomic or protein sequence targets. Since the joined fragments can be separated by non-coding (intronic) regions, their match to the target is termed “spliced alignment”. Even at a genomic level, a brute force application of the spliced alignment approach poses a serious computational burden as it requires aligning each putative fragment to each target as well as recovering the set of putative fragments that best match a target.

Here, we developed MetaEuk, a novel and sensitive reference-based approach to identify single- and multi-exon protein-coding genes in eukaryotic metagenomic data. MetaEuk takes as input a set of assembled contigs and a reference database of target protein sequences or profiles. MetaEuk scans each contig in all six reading frames and extracts putative protein fragments between stop codons in each frame. Thus, MetaEuk makes no assumption about the splicing signal and does not rely on any preceding binning step. MetaEuk uses the MMseqs2 code library [33] for a very fast, yet sensitive identification of putative exons within the fragments. This step also discards the vast majority of fragments, which significantly reduces the computation time of all succeeding steps. The combinatorial task of considering all possible sets of putative exons to best match a given target is solved by means of dynamic programming. Since MetaEuk uses a homology-based strategy to identify protein-coding genes, it can directly confer annotations to its predictions from the matched target proteins.

We benchmarked MetaEuk by using annotated genomes and proteins of seven unicellular organisms from different parts of the eukaryotic tree of life under conditions of increasing evolutionary distance to sequences in the reference database. Despite its high speed and low false positive rates, MetaEuk is able to discover a large fraction of the known proteins in these benchmark genomes. We next applied MetaEuk to study marine eukaryotes. We assembled all Tara Oceans metagenomic samples [20] and focused on ∼1,300,000 contigs of at least 5kbp in length. We clustered more than 330,000,000 proteins to create a comprehensive catalog of over 87,000,000 protein profiles to serve as a reference database. We found the MetaEuk collection of >12,000,000 marine proteins is highly diverged, offering major eukaryotic lineage expansions.

## Results

### The MetaEuk algorithm

The main steps of the algorithm are presented schematically in Figure 1 and a detailed description is provided in the Methods section. For each input contig, all possible protein-coding fragments are translated in six reading frames and searched against a reference target database of protein sequences or profiles. Fragments from the same contig and strand that hit a reference target T are examined together. In each fragment, only the part that was aligned to the target protein T is considered as a putative exon. The putative exons are ordered according to their start position on the contig. Based on their contig locations and the locations of their aligned region on the target T, any two putative exons are either compatible or not. A dynamic programming procedure recovers the highest scoring path of compatible pairs of putative exons by computing the maximum scores of all paths ending with each putative exon. Since homologies among targets in the reference database can lead to multiple calls of the same protein-coding gene, redundancies are reduced by clustering the calls. To that end, all calls are ordered by their start position on the contig. The first call defines a new cluster and all calls that overlap it on the contig are assigned to its cluster if they share an exon with it. The next cluster is defined by the first unassigned call. After all calls are clustered, the best scoring call is selected as the representative of the cluster, termed a “prediction”. Finally, as overlaps of genes on the same strand are very rare [as reviewed by 34], gene predictions overlapping others on the same strand with a better E-value are removed.

**Figure 1.**
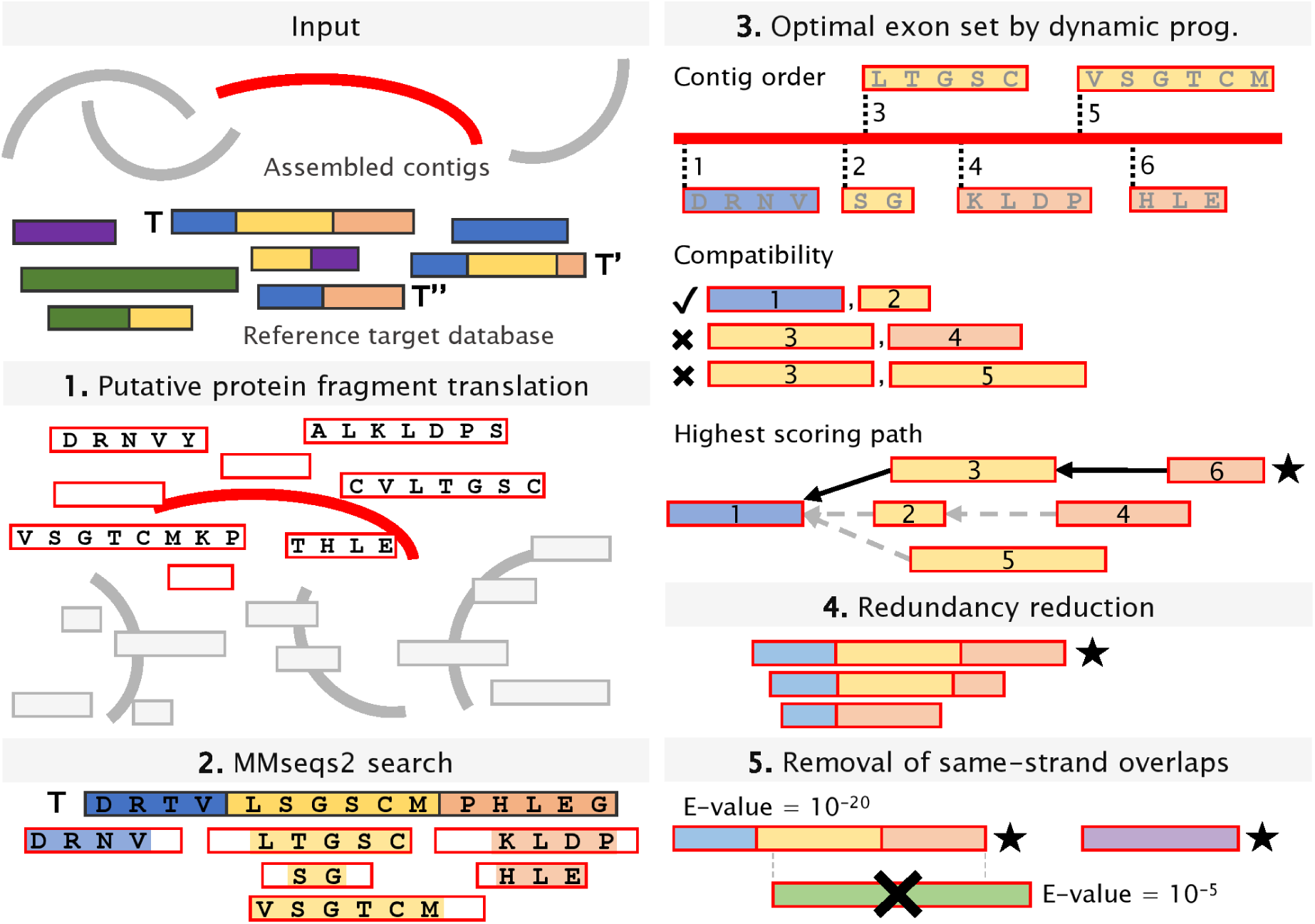
MetaEuk algorithm. Input to MetaEuk are assembled metagenomic contigs and a reference database of protein sequences. (1) Six-frame translation of all putative protein-coding fragments from each contig. (2) Fragments on the same contig and strand that hit the same reference protein T are examined together. (3) Putative exons are identified and ordered according to their start position on the contig. The highest score and path (denoted with a star) of a set of compatible putative exons is computed by dynamic programming, in which individual scores of the putative exons are summed and unmatched amino acids are penalized. (4) Redundancies amongst gene calls due to homologous targets (T, T’ and T’’) are reduced and a representative prediction (denoted with a star) is retained. (5) Contradicting predictions of overlapping genes on the same strand are resolved by excluding the prediction with the higher E-value.

### Performance evaluation on benchmark data

We evaluated MetaEuk using seven annotated unicellular eukaryotic organisms obtained from the NCBI’s genome assembly database [35] (Table 1). These organisms are varied in terms of their phylogenetic group, genome size, number of annotated proteins, fraction of multi-exon genes, and assembly quality. MetaEuk was run on the assembled scaffolds of each of these organisms against the UniRef90 [36] database with an average run time of 42 minutes per genome, or 0.5 Mbp/min, on a server with two 8-core with Intel Xeon E5-2640v3 CPUs and 128 GB RAM (Table 1). The NCBI data included the scaffold coordinates of the annotated protein-coding genes and their exons. In the following sections we used this information to assess MetaEuk’s sensitivity and precision by mapping MetaEuk predictions to annotated proteins in their scaffold location. This was done based on the scaffold boundaries of the MetaEuk prediction and the annotated protein and by requiring high sequence identity of their protein alignment. We then computed the coverage of individual exons of the annotated proteins to which MetaEuk predictions were mapped. These mappings are fully described in the Methods section.

**Table 1.** Species used to benchmark MetaEuk.

| Species | Group | Genome size (Mbp) | # scaffolds | # annotated proteins | % multi-exon proteins | GC% | MetaEuk run time against UniRef90 |
| --- | --- | --- | --- | --- | --- | --- | --- |
| *Schizosaccharomyces pombe* | Fungi | 12.59 | 4 | 5,132 | 47% | 36 | 35m |
| *Acanthamoeba castellanii* str. Neff | Amoebozoa | 42.02 | 384 | 14,974 | 91% | 57.8 | 59m |
| *Phytomonas* sp. isolate EM1 | Excavata | 17.78 | 138 | 6,381 | 0% | 48 | 37m |
| *Babesia bigemina* | Alveolates | 13.84 | 483 | 5,079 | 54% | 50.6 | 35m |
| Nucleomorph of *Lotharella oceanica* | Rhizaria | 0.68 | 4 | 668 | 39% | 32.8 | 24m |
| *Phaeodactylum tricornutum* | Stramenopiles | 27.45 | 88 | 10,408 | 46% | 48.8 | 51m |
| *Aspergillus nidulans* | Eurotiomycetes | 30.28 | 91 | 9,556 | 88% | 50.3 | 52m |

### Sensitivity at evolutionary distance

Sequences from major eukaryotic clades, such as Rhizaria, Stramenopiles, and Dinoflagellata are poorly represented in public protein databases, despite their high abundance in the environment [17]. We therefore measured the ability of MetaEuk to identify homologous protein-coding genes in organisms, which have distant evolutionary relatives in the reference database, as would be the case in a typical metagenomic analysis. To that end, for each annotated organism, we considered five sets of MetaEuk predictions. The first is the base set, which consisted of all predictions. Since we worked with annotated species, their proteins are well represented in UniRef90. The base set therefore reflects ideal conditions, in which the queried organisms are close to the reference database. The other four sets reflect an increasing evolutionary distance and were generated by excluding MetaEuk gene calls whose Smith-Waterman alignment (computed using MMseqs2) to their UniRef90 target had more than 90%, 80%, 60% or 40% sequence identity. We measured sensitivity as the fraction of annotated proteins from the query genome to which a MetaEuk prediction was mapped (see Methods). For all organisms, the sensitivity of the base set of predictions was at least 92%, and sensitivity decreased with the sequence identity threshold (Figure 2A). However, even at low thresholds (40% – 60%), a significant fraction of the annotated proteins were discovered.

**Figure 2.**
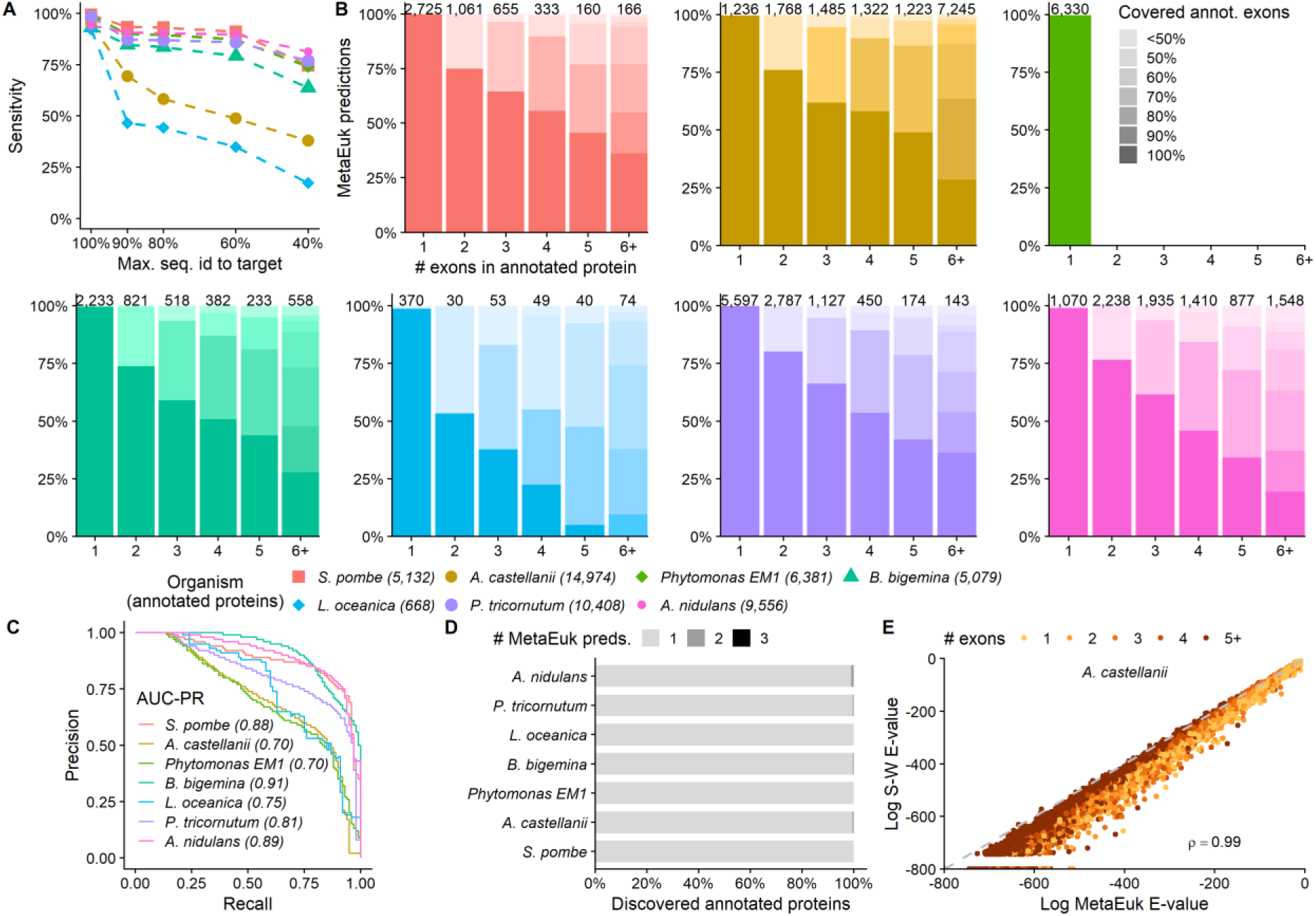
MetaEuk evaluation on benchmark. MetaEuk predictions were mapped to annotated proteins. (A) Conditions of increasing evolutionary divergence were simulated by excluding gene calls based on their sequence identity to their target. Sensitivity is the fraction of annotated proteins from the query genome to which a MetaEuk prediction was mapped. (B) Fraction of exons covered by MetaEuk (color saturation). The number of MetaEuk predictions is indicated on top of each bar. (C) In an annotation-dependent precision estimation MetaEuk predictions that mapped to an annotated protein were considered as *true* and the rest as *false*. These sets of predictions are well separated by their E-values, as indicated by the high AUC-PR values. (D) Fraction of annotated protein-coding genes that were split by MetaEuk into two (dark grey) or three (black) different predictions. (E) Comparison of the E-values computed by MetaEuk and by the Smith-Waterman algorithm for *A. castellani* proteins. The Spearman rho indicates high correlation for *A. castellani* and the other organisms (Supp. Figure 3A).

### Annotated exon coverage

We next assessed MetaEuk’s performance at the level of individual exons. For each MetaEuk prediction from the base set and its mapped annotated protein, we computed the proportion of annotated exons that were covered by the prediction (see Methods). Overall, the majority of predictions covered the majority of exons and, as expected, the fraction of predictions that cover all annotated exons decreases with the number of exons in the annotated protein (Figure 2B). For all organisms, most (77% – 91%) annotated exons were covered by MetaEuk predictions. In addition, we found that the fraction of multi-exon MetaEuk predictions was similar to that presented in Table 1 (average difference: 10%, Supp. Figure 1A) and that single-exon predictions tended to have longer exons than multi-exon predictions (Supp. Figure 1B). An additional measure of completeness of MetaEuk predictions is the coverage of the target UniRef90 protein based on which the prediction was made. We therefore aligned each predicted MetaEuk protein to its target and found that on average, > 83% of predictions covered > 90% of their target (Supp. Figure 2).

### Precision

MetaEuk predictions that were mapped to annotated proteins were considered as true predictions. We first measured the precision of MetaEuk by using the NCBI annotations as gold standard and regarded all predictions in the base set that were not mapped to an annotated protein (8% – 35%, Supp. Figure 2) as false. We computed precision-recall curves by treating the predictions’ E-values as a classifying score. We found good separation (AUC-PR > 0.7 in all cases) between predictions that mapped to annotated proteins and the rest (Figure 2C). However, a prediction that does not map to a known protein is not necessarily false as it might reflect an unannotated protein. We found that about 40% of the unmapped predictions overlap a protein-coding gene on the opposite strand or are on scaffolds that had no annotation at all (Supp. Figure 2), suggestive of post-hoc exclusion criteria in the NCBI annotation procedure. For this reason, we also measured the precision of MetaEuk independently of external annotations by using an inverted-sequence null model. For this annotation-free approach, we ran standard MetaEuk on the inverted sequences of the six frame-translated putative fragments. Each prediction based on these inverted sequences can therefore be considered a false positive. We applied the same E-value cutoff for reporting predictions based on the original sequence data and based on the inverted set. For all organisms, the total number of false positive predictions produced by this approach was low (0 – 12), indicating very high precision (> 99.9%).

### Redundancy reduction

MetaEuk’s redundancy reduction procedure divides gene calls into disjoint clusters and retains a representative call as gene prediction for each cluster (see Methods). This reduces the number of potential protein-coding genes that need to be inspected. E.g., for *S. pombe*, MetaEuk produced over 1,100,000 calls that were reduced to a total of 5,564 predictions in the base set. A full reduction of redundancy is achieved when no two predictions correspond to same protein-coding gene. We thus identified cases in which two or more MetaEuk predictions were mapped to the same protein-coding gene. We found that for all benchmark organisms, redundancy is greatly reduced, as more than 99% of the annotated protein-coding genes in the benchmark scaffolds are only predicted once (Figure 2D).

### Statistical scores

For each prediction, MetaEuk computes a bit-score between the set of translated and joined putative exons and the target protein. Based on this bit-score and the size of the reference database, an E-value is computed (see Methods). We evaluated MetaEuk’s bit-scores and E-values by comparing them to those computed for each predicted protein and its target by the Smith-Waterman algorithm. Since MetaEuk penalizes missing and overlapping amino acids when joining putative exons, we expect the MetaEuk bit-score to be more conservative than the direct Smith-Waterman alignment bit-score. We found very high levels of agreement between the MetaEuk statistics and the Smith-Waterman statistics (Figure 2E, Supp. Figure 3). This suggests a straightforward statistical interpretation of MetaEuk prediction scores.

### Effect of contig length

Assembling metagenomic reads often produces contigs that are much shorter than the scaffolds of the organisms we used for benchmarking MetaEuk (Table 1). We thus aimed to assess the effect of analyzing shorter genomic stretches by artificially dividing each of the scaffolds from Table 1 into shorter contigs following a typical length distribution with a minimum of 5kbp in length and a median of 6.8kbp (see Methods). Any protein-coding gene that spans more than one contig is expected to result in incomplete MetaEuk predictions. Indeed, while the sensitivity measured by the mapping to annotated proteins remained similar to that recorded on the original scaffolds (Supp. Figure 4A), we found that more predictions were partial and covered fewer annotated exons (Supp. Figure 4B) as well as an increase of up to 15% in annotated genes being split into several MetaEuk predictions (Supp. Figure 4D).

### Eukaryotic protein-coding genes in the ocean

To date, little is known about the biological activities of eukaryotes in the oceans [2, 37]. We aimed to use MetaEuk to discover eukaryotic protein-coding genes in the Tara Oceans metagenomic dataset [20]. We first used MEGAHIT [38] to assemble all 912 samples of this project. We retained 1,351,204 contigs of at least 5kbp in length that were classified as potentially eukaryotic by EukRep [27]. We next constructed a comprehensive set of reference proteins by uniting over 21,000,000 representative sequences of the Uniclust50 database [39], the MERC dataset of over 292,000,000 protein sequence fragments assembled from eukaryotic Tara Oceans metatranscriptomic datasets [40], and over 18,500,000 protein sequences of MMETSP, the Marine Microbial Eukaryotic Transcriptome Sequencing Project [17, 41]. We clustered the joint dataset of 331,913,793 proteins using the combined Linclust / MMseqs2 four-step cascaded clustering workflow [42] with a minimal sequence identity of 20% and high sensitivity (-s 7). This resulted in 87,984,812 clusters, most of which (> 97%) contained proteins from a single reference dataset (Figure 3). For each cluster, a multiple sequence alignment was generated, based on which a sequence profile was computed.

**Figure 3.**
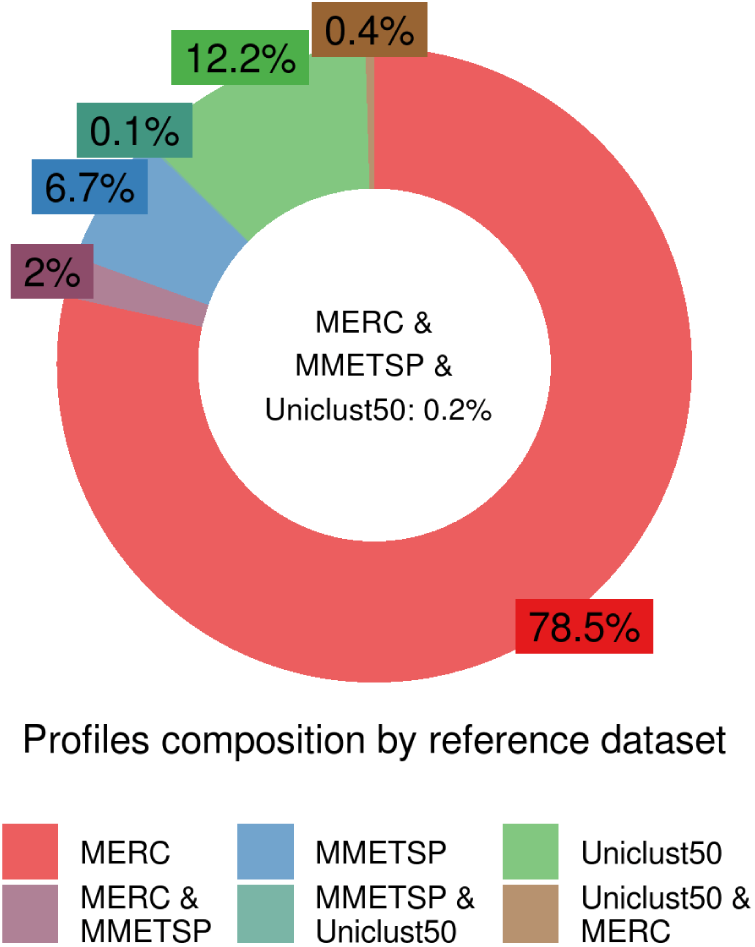
Reference profiles composition. Proteins from three datasets: MERC (292 million), MMETSP (18.5 million) and Uniclust50 (21 million) were clustered into ∼88 million clusters. Most clusters contained proteins from a single reference dataset. The profiles computed based on these clusters served as the reference database for the MetaEuk run on the Tara Oceans contigs.

MetaEuk’s run using this reference database took eight days on ten 2×8-core servers and resulted in 12,111,301 predictions with no same-strand overlaps in 1,287,197 of the Tara Oceans contigs. Due to sequence similarities among the assembled contigs, some of these proteins are identical to each other, leaving a total of 6,158,526 unique proteins. We examined the distribution of predictions per contig, the number of putative exons in each prediction and the length of putative exons in single-exon and multi-exon predictions. We found that the number of predictions increases as a function of the contig length (Figure 4A), about 24% of predictions had more than one putative exon (Figure 4B) and multi-exon predictions tend to have shorter putative exons than single-exon predictions (4C). We analyzed the contribution of each reference dataset to the profiles based on which the MetaEuk predictions were made. MERC, MMETSP and Uniclust50 contributed 77.4%, 5.7% and 4.3% of the predictions, respectively. The rest of the predictions were based on mixed-dataset clusters (Supp. Figure 5). We then used MMseqs2 to query the MetaEuk predicted proteins against their targets. Over 33% of the MetaEuk predictions have less than 60% sequence identity to their MERC, MMETSP or Uniclust50 target (Figure 5A). Finally, we found that 70% of the MetaEuk predicted proteins covered at least 80% of their reference target (Figure 5B).

**Figure 4.**
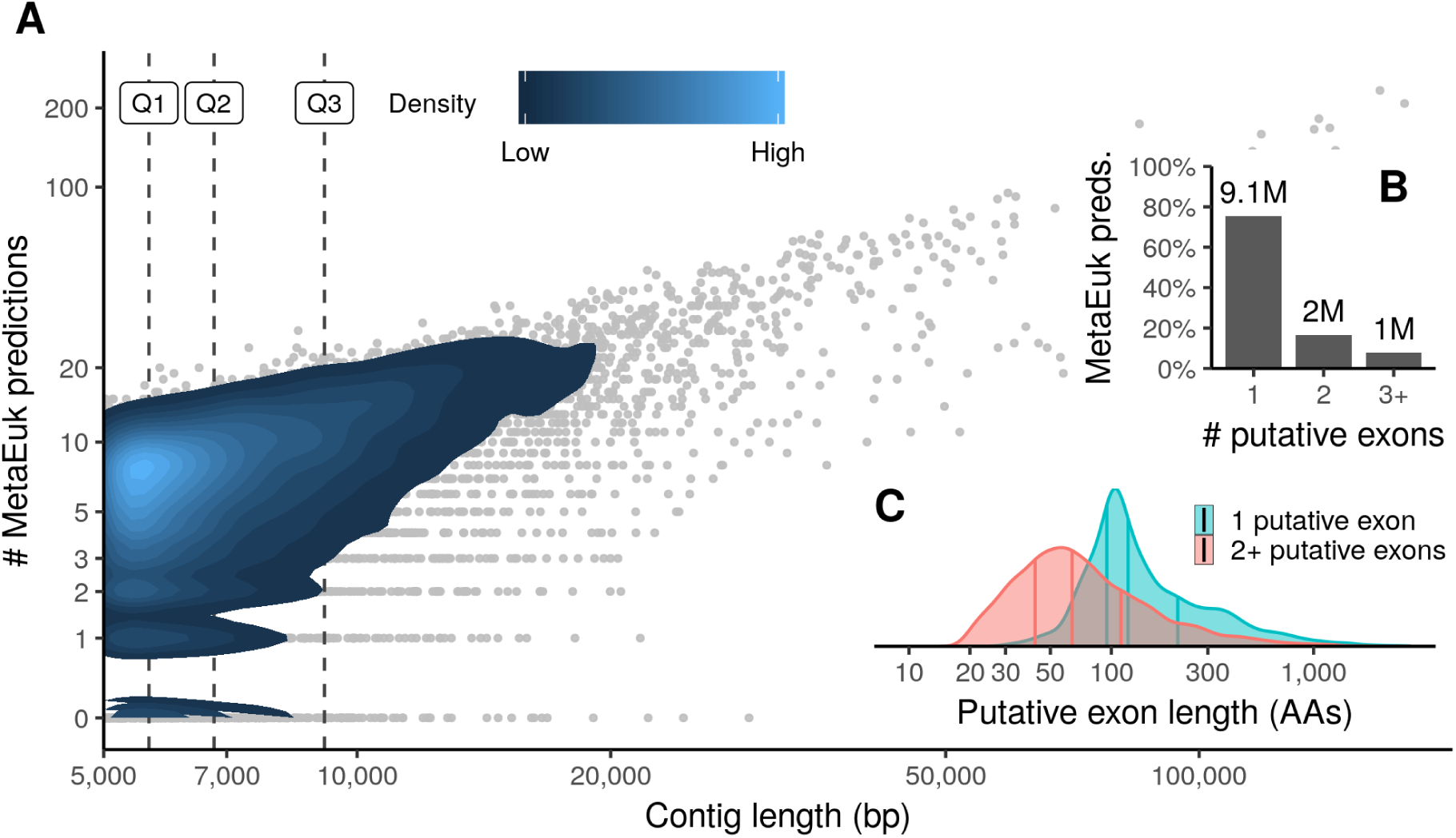
MetaEuk predictions on Tara Oceans contigs. MetaEuk was run on over 1.3 million contigs assembled from Tara Oceans metagenomic reads against a reference database of ∼88 million protein profiles. (A) The number of MetaEuk predictions per contig increases with its length. Horizontal lines mark contig length quartiles. (B) Most (76%) MetaEuk predictions had a single putative exon. The absolute number of predictions is indicated above each bar. (C) Single-exon predictions tend to have longer putative exons than multi-exon predictions.

**Figure 5.**
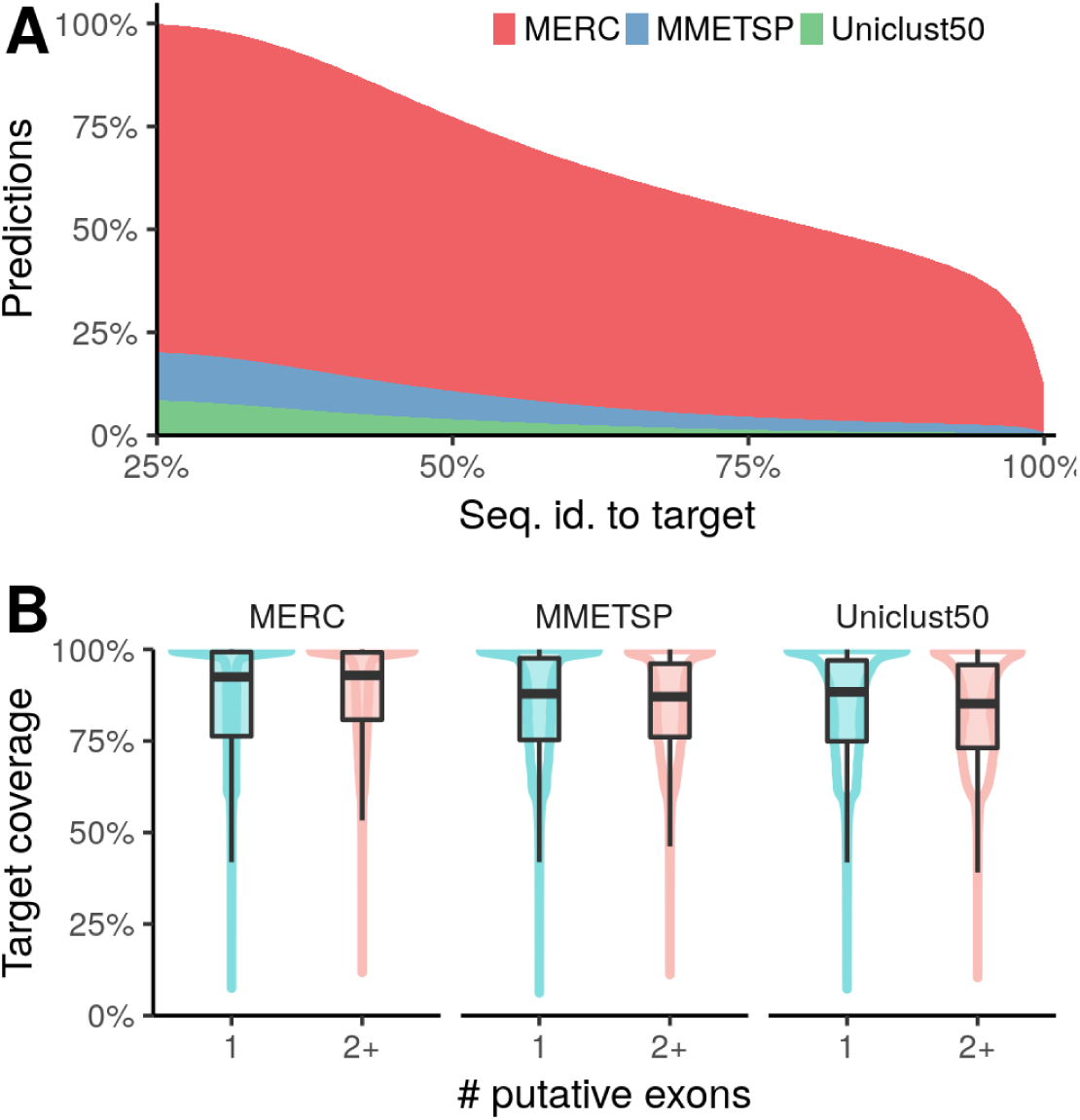
MetaEuk predictions compared to the reference datasets. MetaEuk predicted proteins were queried against the representative sequence of their target reference cluster. (A) About one third of the predicted MetaEuk proteins had less than 60% sequence identity to their target. (B) Targets are well covered by MetaEuk predicted proteins.

We next explored the taxonomic composition of the MetaEuk proteins. Since the majority (77%) of MetaEuk predictions were based on homologies to the MERC dataset, for which no taxonomic annotation is available, we queried the MetaEuk marine proteins collection against the Uniclust90 dataset [39] and the MMETSP dataset, both annotated using NCBI taxonomy (see Methods). We found that 63% of predictions based on homologies to the MERC dataset did not match any protein in either of the reference datasets, which means ∼49% (63% of 77%) of the MetaEuk marine proteins collection could not be assigned any taxonomy. This is in agreement with 52% of unassigned unigenes assembled from Tara Oceans metatranscriptomics [20]. We next assigned taxonomic labels to each assembled contig by conferring the taxonomic label with the best E-value of all MetaEuk predictions in the contig. This allowed us to annotate 92% of the contigs for which MetaEuk produced predictions (87% of all input contigs). We found that 82% of the contigs were assigned to the domain Eukaryota and 9% to non-eukaryotes, mostly bacteria (Figure 6A). We then examined the assigned eukaryotic supergroups below the domain level. About 12% of the eukaryotic contigs could not be assigned a supergroup. Among the most abundant eukaryotic supergroups are Metazoa and Chlorophyta (Figure 6B).

**Figure 6.**
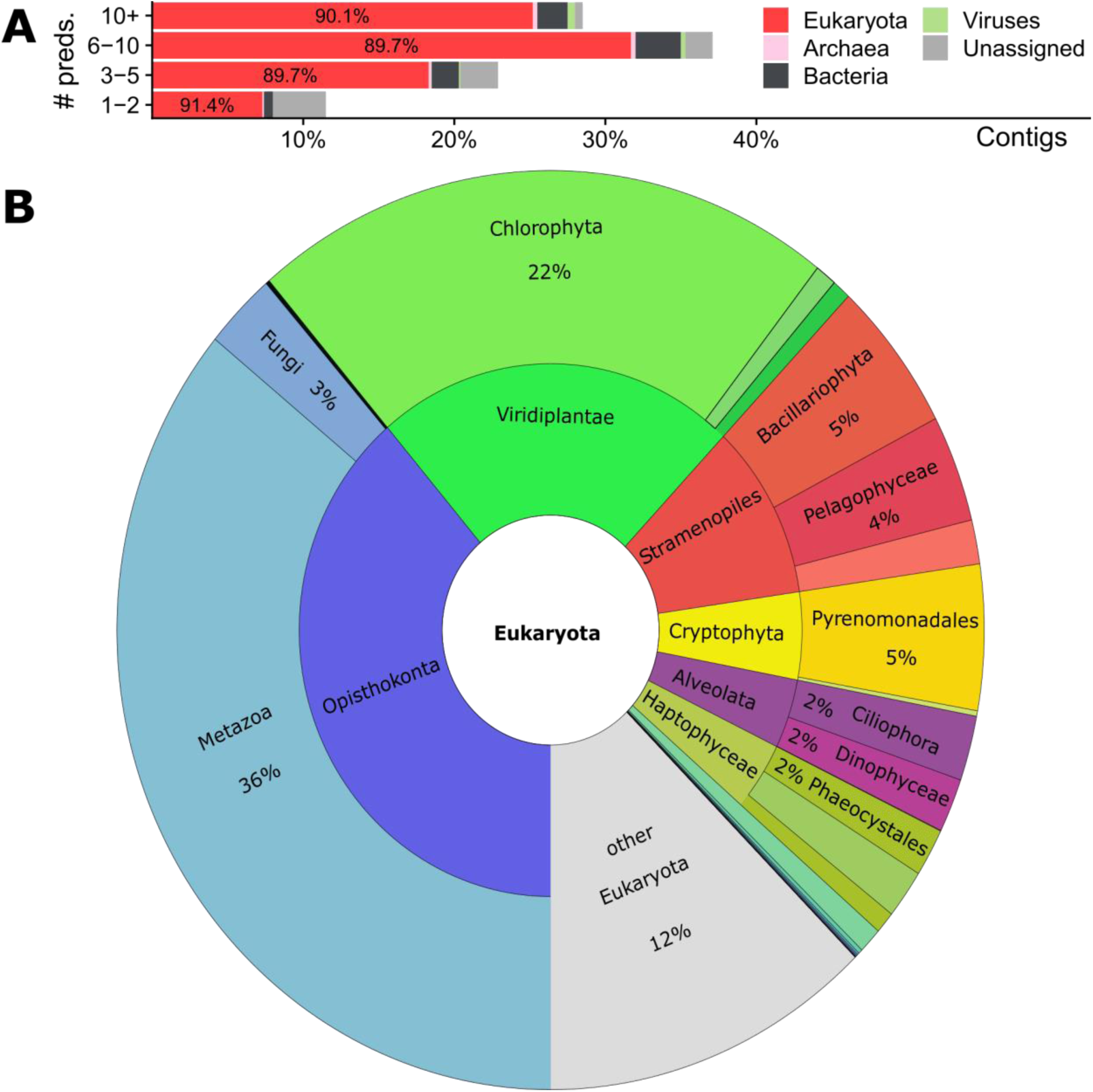
Taxonomy of Tara Oceans contigs with MetaEuk predictions. The best-scoring taxonomic label of all predictions on each contig was conferred to the contig. Contigs were divided into four categories according to their number of MetaEuk predictions. Over 82% of the contigs were assigned to the domain Eukaryota. (A) The proportion of unassigned contigs decreases with the number of MetaEuk predictions on the contig. The fraction of eukaryotic contigs out of all assigned contigs is about 90% in all four categories. (B) Eukaryotic taxonomic labels below the domain level.

The high fraction of unassignable predictions (49%) prompted us to seek an additional way to assess the diversity of the MetaEuk marine proteins. We thus collected orthologous sequences of the large subunits of RNA polymerases, which are universal phylogenetic markers [43] from 985 organisms for which we had taxonomic information, as well as 1,076 MetaEuk proteins, which consisted of all five Pfam domains of the large subunit in the right order (see Methods). We aligned these sequences using MAFFT [44] and constructed the maximum-likelihood phylogeny using RAxML [45]. The aim of this analysis was to delineate the diversity of eukaryotic taxa of the MetaEuk marine proteins collection and not to resolve the exact phylogenetic relationships among them. As can be seen in Figure 7, MetaEuk proteins offer major lineage expansions in under-sampled eukaryotic supergroups. Importantly, the strict ortholog collection procedure performed for this analysis results in a conservative estimate of the diversity level of the MetaEuk marine proteins collection.

**Figure 7.**
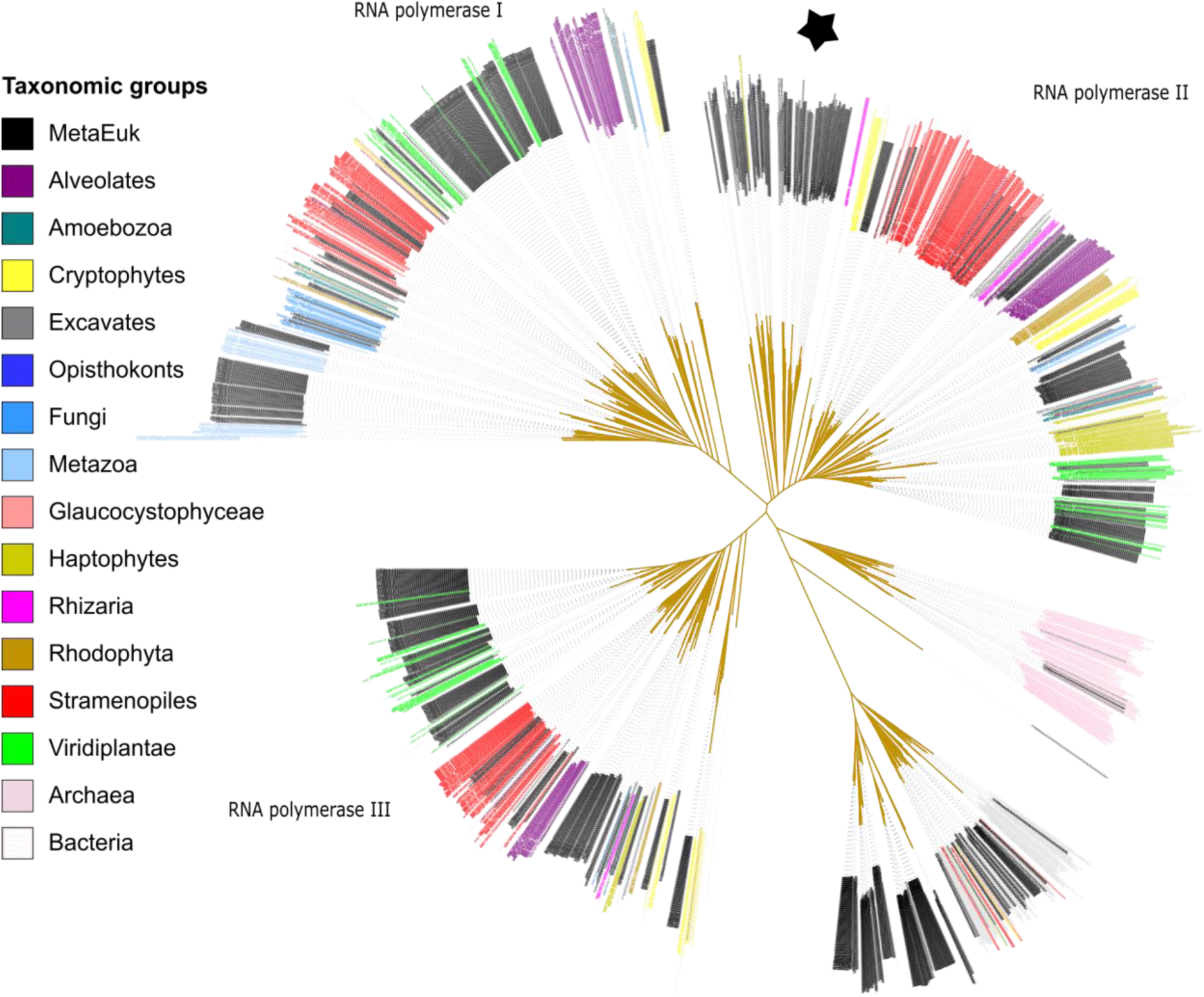
Diversity of MetaEuk marine eukaryotic proteins. Homologous sequences of the large subunits of RNA polymerases of 985 species as well as 1,076 MetaEuk marine proteins were collected and a maximum-likelihood tree was computed based on their alignment. MetaEuk sequences (black) expand major eukaryotic lineages, including deeply rooted supergroups (denoted with star).

## Discussion

We presented MetaEuk, an algorithm designed for large-scale analysis of eukaryotic metagenomic data. We demonstrated its utility for discovering proteins from highly diverged eukaryotic groups by analyzing assemblies of a huge set of 912 marine metagenomics samples. MetaEuk makes no assumption concerning splice site signatures and does not require a preceding binning procedure, which renders it suitable for the analysis of contigs from a mixture of highly diverged organisms. In order to achieve this, MetaEuk considers all possible putative protein-coding fragments from each input contig. Applying the spliced alignment dynamic programming procedure to recover the optimal set of putative exons directly on these fragments would result in a run time complexity per contig that is quadratic in the number of its fragments times the number of targets in the reference database. This is not feasible for metagenomics, as the number of fragments can be hundreds of millions (e.g., from 1,351,204 Tara Oceans contigs, 152,519,258 fragments were extracted) and the reference database should be as comprehensive as possible (in this study, we used more than 87,000,000 protein profiles). To circumvent this limitation, MetaEuk takes advantage of the ultra-fast MMseqs2 search algorithm, which allows it to find putative exons matching a reference protein sequence with sufficient significance (in this study, a lenient E-value of 100). MetaEuk does not require significance at the exon level as it can combine sub-significant single exon matches to highly significant multi-exon matches. For example, two putative exons each with an E-value of 10 (corresponding to a bit-score of 25-40 in this study), are not individually significant but the sum of their bit-scores of at least 50 corresponds to a significant E-value of 1E-05.

MetaEuk is not designed to recover accurate splice sites, but rather to identify the protein-coding parts within exons. Indeed, we showed that MetaEuk predictions on the benchmark covered the majority (77% – 91%) of exons in annotated proteins. Since MetaEuk relies on local alignment at the amino acid level, it could potentially report pseudogenes, which still bear sequence similarity to reference proteins. However, we found that the majority of benchmark predictions (65% – 92%) mapped to NCBI annotated protein-coding genes, while the rest could be well separated from those that mapped by their E-values (AUC-PR > 0.7). Furthermore, unmapped predictions can reflect a missing annotation or post-hoc exclusion criteria (e.g., removal of annotations that overlap a better scoring one on the opposite strand). We therefore measured precision independently of annotations by running standard MetaEuk on the inverted sequences of the putative protein fragments extracted from the contigs. By using this annotation-free approach, we showed that MetaEuk’s precision was greater than 99.9% for all benchmark organisms. Put together, MetaEuk’s strength is in describing the protein-coding repertoire of versatile environments rather than in constructing statistical models of exon-intron transitions.

The Tara Oceans contigs analyzed in this study were assembled from Illumina HiSeq 2000 short reads. High population diversity, repeat regions, and sequencing errors are among the major factors contributing to the computational challenge associated with metagenomic assembly [reviewed by 46]. These factors reduce the quality of the assembly as reflected, for example, in shorter contig lengths, chimeric contigs and contigs containing strand inversions. These in turn, directly and negatively impact MetaEuk. Shorter contigs limit its ability to discover multi-exon protein-coding genes as it searches for them within a contig. In addition, predictions on contig edges can be partial, which is more likely to happen in a highly fragmented assembly. By dividing each of the benchmark scaffolds to contigs whose lengths were drawn at random based on the length distribution of the Tara Ocean contigs, we showed that while MetaEuk retains its overall sensitivity to detect protein coding genes even under conditions of increasing evolutionary distance between the query organism and the target reference database, the completeness of its predictions is reduced. We thus expect MetaEuk to benefit from future improvements in assembly algorithms, higher sequencing coverage, and long-read sequencing technology [47–50].

In addition to developing MetaEuk, we generated two useful resources for the analysis of eukaryotes as part of this study. The first is the comprehensive protein profile database, which was computed using protein sequences from three sources: MERC, MMETSP and Uniclust50. With ∼88 million records, it is the largest profile database focused on eukaryotes to date. Since MERC was assembled from the Tara Oceans metatranscriptomic data, we expected it to be a valuable resource for discovering protein-coding genes in the same environment. Indeed, we found that the majority of MetaEuk predictions (77%) were based on MERC protein profiles. Furthermore, the high fraction of MERC-based predictions that could not be assigned a taxonomic label (63%) demonstrates the uniqueness of this resource.

The second resource is the MetaEuk marine protein collection, which is available on our search webserver (https://search.mmseqs.com/search) for easy investigation [51]. Using a phylogenetic marker protein, we showed that this collection contains proteins spanning major eukaryotic lineages, including supergroups with very few available genomes. Over 33% of these proteins have less than 60% sequence identity to the representative reference proteins that were used to predict them, indicating their diversity with respect to the reference database. Unlike the MERC and MMETSP proteins, MetaEuk proteins are predicted in the context of genomic contigs. This allows us to learn of the number of putative exons that code for them as well as to examine them together with other proteins on the same contig. The latter is useful for conferring taxonomic annotations to unlabeled predictions on the same contig as well as for detecting complex functional modules, by searching for co-occurrences of the module’s proteins on the same contig.

As was demonstrated by the challenge of assigning taxonomy to highly diverged eukaryotic proteins, the paucity of eukaryotic sequences in reference databases is currently a major limitation in the study of eukaryotes. Thus, we expect the resources produced in this study and further analyses of eukaryotic metagenomic data using MetaEuk to produce a more comprehensive description of the tree of life [16,52–54].

## Conclusions

MetaEuk is a sensitive reference-based algorithm for large-scale discovery of protein-coding genes in eukaryotic metagenomic data. Applying MetaEuk to large metagenomic datasets is expected to significantly enrich our databases with highly diverged eukaryotic protein-coding genes. By adding sequences from under-sampled eukaryotic lineages, we can improve sequence homology searches, protein profile computation and thereby homology-based function annotation, template-based and even de-novo protein structure prediction [55, 56]. These, in turn will allow for further exploration of eukaryotic activity in various environments [57].

## Methods

### MetaEuk algorithm

#### Code and resources availability

The MetaEuk source code, compilation instructions and a brief user guide are available from https://github.com/soedinglab/metaeuk under the GNU General Public License v3.0. The resources produced during this study are available from http://wwwuser.gwdg.de/~compbiol/metaeuk/.

#### Putative exons compatibility

In the first two stages of the MetaEuk algorithm all possibly coding protein fragments are translated from the input contigs. We scan each contig in six frames and extract the fragments between stop codons. These fragments are queried against the reference target database using MMseqs2. A set of fragments from the same contig and strand that have local matches to the same specific target *T* define a set of putative exons. We say two putative exons *P*_*i*_ and *P*_*j*_ from the same set are compatible with each other if they can be joined together to a multi-exon protein.

Each *P*_*i*_ is associated with four coordinates: the amino-acid position on *T* from which the match to *P*_*i*_ starts 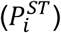 and ends 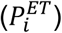; the nucleotide position on the contig from which the translation of *P*_*i*_ starts 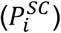 and ends 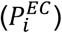. We require a match of at least 10 amino acids (a minimal exon length). We consider putative exons *P*_*i*_ and *P*_*j*_ with 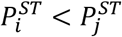 as compatible on the plus strand if:

(1) their order on the contig is the same as on the target: 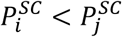;
(2) the distance between them on the contig is at least the length of a minimal intron but not more than the length of a maximal intron: 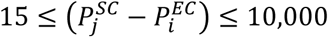;
(3) their matches to *T* should not overlap. In practice we allow for a short overlap to account for alignment errors: 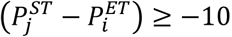.

In case *P*_*i*_ and *P*_*j*_ are on the negative strand, we modify conditions (1) and (2) accordingly:

(1) 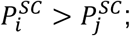
(2) 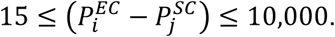

Since the adjustment of conditions to the minus strand is straightforward, in the interest of brevity we focus solely on the plus strand in the following text.

We say a set of *k* > 1 putative exons is compatible if, when ordered by their 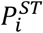 values, each pair of consecutive putative exons is compatible. (A set of a single exon is always compatible).

### Bit-score and E-value computation

A set of *k* compatible putative exons defines a pairwise protein alignment to the target *T*. This alignment is the concatenation of the ordered local alignments of all putative exons to *T*. Between each consecutive putative pair of exons *P*_*i*_ and *P*_*i*+1_ there might be unmatched amino acids in *T* or there might be a short overlap of their matches to *T*. We denote the number of unmatched amino acids between *P*_*i*_ and *P*_*i*+1_ as *l*_*i*_, which can take a negative value in case of an overlap. MetaEuk computes the bit-score of the concatenated pairwise alignment *S*(*P*_*set*_, *T*) by summing the individual Karlin-Altschul [58] bit-scores *S*(*P*_*i*_,*T*) of the putative exons to *T* and penalizing for unmatched or overlapping amino acids in *T* as follows:

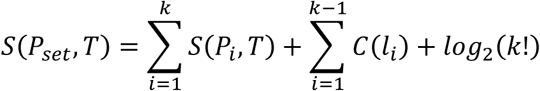

where the penalty function is *C*(*l*_*i*_) = −|*l*_*i*_| for *l*_*i*_ ≠ 1 and 0 if *l*_*i*_ = 1. The last term rewards the correct ordering of the *k* exons.

An E-value is the expected number of matches above a given bit-score threshold. Since for each contig, at most one gene call is reported per strand and target in the reference database, the E-value takes into account the number of amino acids in the reference database *D* and the search on two strands:

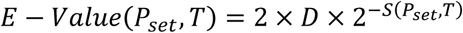

### Dynamic programming

Given a set of *n* putative exons and their target, MetaEuk finds the set of compatible exons with the highest combined bit-score. First, all putative exons are sorted by their start on the contig, such that 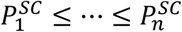. The dynamic programming computation iteratively computes vectors *S*, *k*, and *b* from their first entry 1 to their *n*^*t*ℎ^. The entry *S*_*i*_ holds the score of the best exon alignment ending in exon *i* and *k*_*i*_ holds the number of exons in that set. Once the maximum score is found, the exon alignment is back traced using *b*, in which entry *b*_*i*_ holds the index of the aligned exon preceding exon *i* (0 if *i* is the first aligned exon). Using the following values:

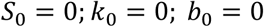

all putative exons *P*_*j*_ are examined according to their order and the score vector is updated:

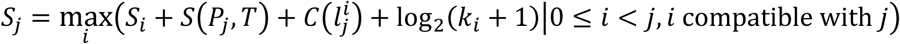

*k*_*j*_ and *b*_*j*_ are updated accordingly. The terms log_2_(*k*_*i*_ + 1) add up to the score contribution 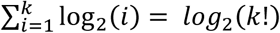 and the transition 0 to *j* is defined as compatible with 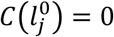 for all *j*. The optimal exon set is then recovered by tracing back from the exon with the maximal score. This dynamic programming procedure has time complexity of *O*(*n*^2^).

### Clustering gene calls to reduce redundancy

MetaEuk assigns a unique identifier to each extracted putative protein fragment (stage 1 in Figure 1). A MetaEuk exon refers to the part of a fragment that matched some target T (stage 2 in Figure 1, tinted background) and has the same identifier as the fragment. Two calls that have the same exon identifier in their exon set are said to share an exon. MetaEuk reduces redundancy by clustering calls that share an exon (stage 4 in Figure 1) and selecting a representative call as the gene prediction of each cluster. To that end, all *N* MetaEuk calls from the same contig and strand combination are ordered according to the contig start position of their first exon. Since this order can include equalities, they are sub-ordered by decreasing number of exons. The first cluster is defined by the first call, which serves as its tentative representative. Let *m* be the last contig position of the last exon of this representative. Each of the following calls is examined so long as its start position is smaller than *m* (i.e., it overlaps the representative on the contig). If that call shares an exon with the representative, it is assigned to its cluster. In the next iteration, the first unassigned call serves as representative for a new cluster and the following calls are examined in a similar manner, adding unassigned calls to the cluster in case they share an exon with the representative. The clustering ends with the assignment of all calls. At this stage, the final prediction is the call with the highest score in each cluster. This greedy approach has time complexity of *O*(*N* × *log*(*N*) + *N* × *A*), where *A* is the average number of calls that overlap each representative on the contig. Since in practice, *A* ≪ *N*, the expected time complexity is *O*(*N* × *log*(*N*)).

### Resolving same-strand overlapping predictions

After the redundancy reduction step, MetaEuk sorts all predictions on the same contig and strand according to their E-value. It examines the sorted list and retains predictions only if they do not overlap any preceding predictions on the list.

### Benchmark datasets

The UniRef90 database was obtained in March 2018. The annotated information of *Schizosaccharomyces pombe* (GCA_000002945.2), *Acanthamoeba castellanii str. Neff* (GCA_000313135.1), *Babesia bigemina* (GCA_000981445.1), *Phytomonas sp. isolate* EM1 (GCA_000582765.1), Nocleomorph of *Lotharella oceanica* (GCA_000698435.2), *Phaeodactylum tricornutum* (GCA_000150955.2), and *Aspergillus nidulans* (GCA_000149205.2) were downloaded from the NCBI genome assembly database (March – September 2018). This information included the genomic scaffolds, annotated protein sequences, and GFF3 files containing information about the locations of annotated proteins and other genomic elements. MetaEuk (Github commit 47141068c171fcdd3d93411ac50958da0f2c4025, MMseqs2 submodule version ebb16f3631d320680a306c03aa7412c572f72ee3) was run with the following parameters: -e 100 (a lenient maximal E-value of a putative exon against a target protein), --metaeuk-eval 0.0001 (a stricter maximal cutoff for the MetaEuk E-value after joining exons into a gene call), --metaeuk-tcov 0.6 (a minimal cutoff for the ratio between the MetaEuk protein and the target) and --min-length 20, requiring putative exon fragments of at least 20 codons and default MMseqs2 search parameters.

### Mapping benchmark predictions to annotated proteins

For each annotated protein, we listed the contig start and end coordinates of the coding part (CDS) of each of its exons. The lowest and highest of these coordinates were considered as the boundaries of the annotated protein, and the stretch between them as its “global” contig length. Similarly, we listed these coordinates and computed the boundaries and global contig length for each MetaEuk prediction. A MetaEuk prediction was globally mapped to an annotated protein if the overlap computed based on their boundaries was at least 80% of the global contig length of either of them and if, in addition, the alignment of their protein sequences mainly consisted of identical amino acids or gaps (i.e., less than 10% mismatches). These criteria allow mapping MetaEuk predictions to an annotated protein, even if they miss some of its exons. Next, we computed the exon level mapping for all globally mapped pairs of MetaEuk predictions and annotated proteins. To that end, we compared their lists of exon contig coordinates. If an exon predicted by MetaEuk covered at least 80% of the contig length of an annotated protein’s exon, we considered the annotated exon as “covered” by the MetaEuk prediction.

### Generating typical metagenomic contig lengths

In order to evaluate MetaEuk’s performance on contigs with a length distribution typical for assemblies from metagenomic samples, we recorded the lengths of the assembled contigs used for the analysis described in the “Tara Oceans dataset” section. The 1,351,204 contigs had a minimal length of 5,002 bps, 1^st^ quartile of 5,661 bps, median of 6,763 bps, 3^rd^ quartile of 9,020 bps and a maximal length of 1,524,677 bps. We divided each annotated scaffold into contigs of lengths that were randomly sampled from these recorded lengths. This resulted in 1,392, 5,061, 1,816, 2,095, 80, 3,153 and 3,273 contigs for *S. pombe*, *A. castellanii*, *Phytomonas sp. isolate* EM1, nucleomorph of *L. oceanica*, *P. tricornutum*, and *A. nidulans*, respectively. MetaEuk was run on these contigs in the same way as on the original scaffolds. Since each of the new contigs corresponded to specific locations on the original scaffolds, we could carry out all benchmark assessments, which relied on mapping between MetaEuk predictions and annotated proteins.

### Tara Oceans dataset

The 912 metagenomic SRA experiments associated with accession number PRJEB4352 were downloaded from the SRA (August – September 2018). The reads of each experiment were trimmed to remove adapters and low quality sequences using trimmomatic-0.38 [59] with parameters ILLUMINACLIP:TruSeq3-PE.fa:2:30:10 LEADING:3 TRAILING:3 SLIDINGWINDOW:4:15 MINLEN:36 (SE for single-end samples). The resulting reads were then assembled with MEGAHIT [38] with default parameters. Contigs of at least 5kbp in length were classified as eukaryotic/non-eukaryotic using EukRep [27], which is trained to be highly sensitive to detecting eukaryotic contigs. MetaEuk was run on the contigs classified as eukaryotic with parameters: -e 100, --metaeuk-eval 0.0001, --min-ungapped-score 35, --min-exon-aa 20, --metaeuk-tcov 0.6, --min-length 40, --slice-search (profile mode) and default MMseqs2 search parameters.

### Taxonomic assignment to predictions and contigs

We used MMseqs2 to query the MetaEuk marine proteins collection against two taxonomically annotated datasets: Uniclust90 and the MMETSP protein dataset. Taxonomic labels associated with each of the MMETSP identifiers were downloaded from the NCBI website (BioProject PRJNA231566). We retained the hit with the highest bit-score value for each prediction if it had an E-value smaller than 1E-05. In addition, we examined the sequence identity between the MetaEuk prediction and the target in order to determine the rank of the taxonomic assignment. Similarly to [20], we used the following sequence identity cutoffs: >95% (species), >80% (genus), >65% (family), >50% (order), >40% (class), >30% (phylum), >20% (kingdom). Lower values were assigned at the domain level. The predictions on each contig were examined and the best-scoring one was used to confer taxonomic annotation to that contig. The assignment was visualized using Krona [60].

### Phylogenetic tree reconstruction

We constructed the tree using the large subunit of RNA polymerases as a universal marker. This subunit contains five RNA_pol_Rpb domains (Pfam IDs: pf04997, pf00623, pf04983, pf05000, pf04998). As detailed below, protein sequences that contained all five domains in the right order were obtained in January-November 2019 from six sources to construct the multiple sequence alignment and tree. The sources were: (1) 75 sequences of the OrthoMCL [61] group OG5_127924. The four-letter taxonomic codes of these sequences were converted to NCBI scientific names, based on information from the OrthoMCL website (http://orthomcl.org/orthomcl/getDataSummary.do). (2) 36 reviewed eukaryotic sequences were downloaded from UniProt [36]. These were used to distinguish between eukaryotic RNA Polymerase I (8 sequences), eukaryotic RNA Polymerase II (16 sequences) and eukaryotic RNA Polymerase III (12 sequences). We then ran an MMseqs2 profile search against the Pfam database (with parameters: -k 5, -s 7) with several query sets and retained results in which all five domains were matched in the right order with a maximal E-value of 0.0001. This allowed us to add the following sources: (3) 674 MMETSP proteins. (4) 100 archaeal proteins; (5) 100 bacterial proteins. For datasets (4) and (5), we first downloaded candidate proteins from the UniProt database by searching for the five domains and restricting taxonomy:archaea (bacteria). We then ran the previously described search procedure and randomly sampled exactly 100 proteins from each group that matched the criterion. (6) 1,076 MetaEuk predictions. The joint set of 2,061 sequences was aligned using MAFFT v7.407 [44] and a phylogenetic tree was reconstructed by running RAxML v8 [45]. Tree visualization was performed in iTOL [62].

## Declarations

### Ethics approval and consent to participate

Not applicable

### Consent for publication

Not applicable

### Availability of data and material

The datasets generated and/or analyzed during the current study are available in http://wwwuser.gwdg.de/~compbiol/metaeuk/

### Authors’ contributions

ELK and JS have designed the MetaEuk algorithm, benchmark and biological application. ELK and MM have developed the algorithm. ELK has analyzed the benchmark and Tara Oceans data. ELK and MM have generated the figures. ELK, JS and MM have drafted the manuscript.

## Acknowledgements

We thank Dr. David Burstein from Tel Aviv University for his helpful insights concerning the phylogenetic analyses and Dr. Christian Woehle from Christian-Albrechts University Kiel for his comments on the manuscript.

