## Supplemental Figures for "MetaEuk – sensitive, high-throughput gene discovery and annotation for large-scale eukaryotic metagenomics"

Running title: Metagenomics eukaryotic gene discovery
Keywords: MetaEuk, eukaryotes, homology detection, prediction, annotation, contigs, marine;

### Supplementary Figures

**Supplementary Figure 1 – MetaEuk predictions by number of exons and exons length**

MetaEuk was run on a benchmark of seven eukaryotic unicellular organisms. (A) The fraction of multi-exon MetaEuk predictions is similar to the fraction of annotated multi-exon proteins (Table 1). (B) Single-exon predictions tend to have longer putative exons than multi-exon predictions.

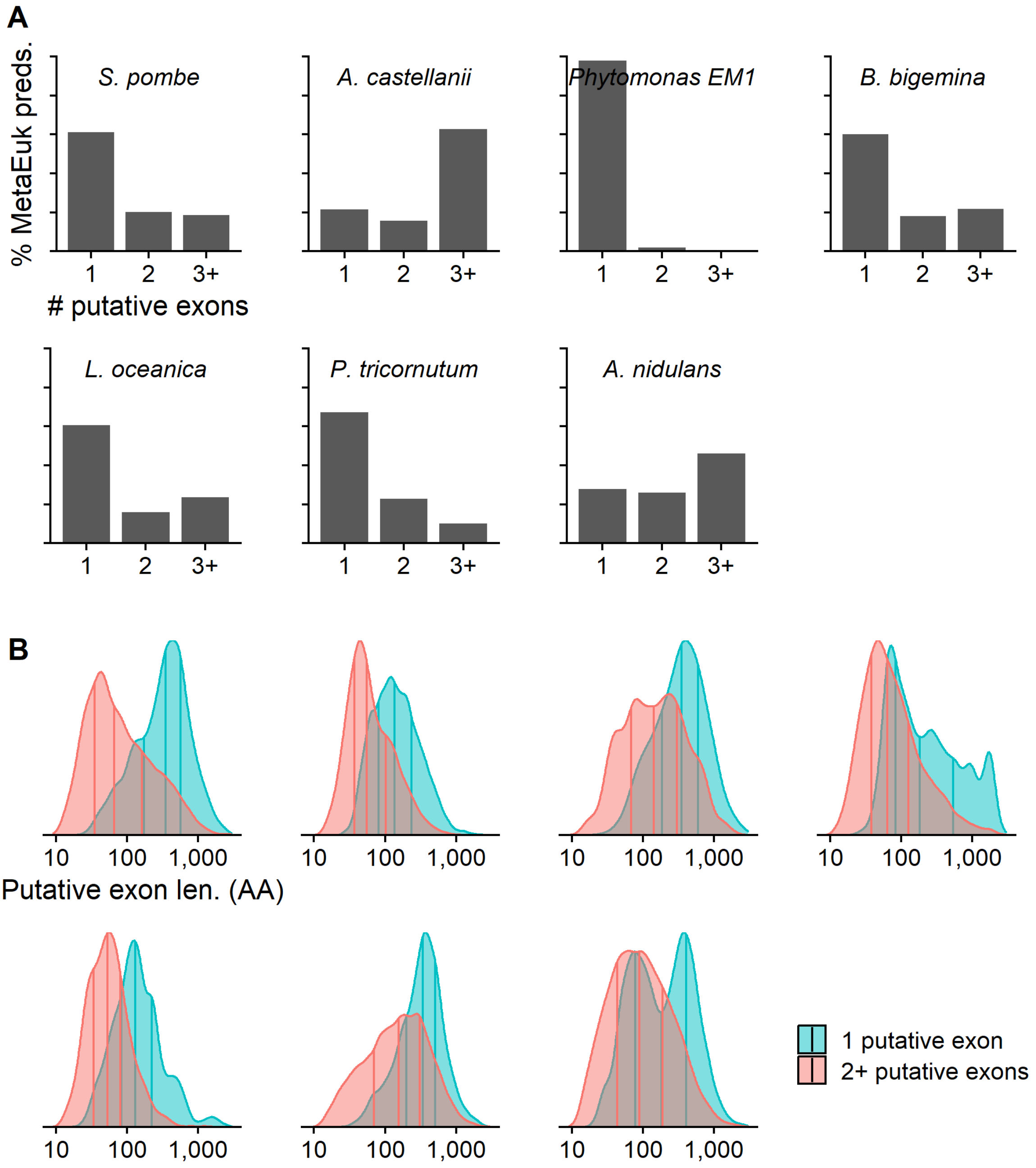

**Supplementary Figure 2 – MetaEuk target coverage**

The protein sequence of each MetaEuk prediction was aligned to the UniRef90 target, which was used to produce the prediction. The level of target coverage was measured while recording the mapping status of the MetaEuk prediction with respect to the annotations of the benchmark organism: mapped to an annotated protein ("prot"), overlap of at least ten nucleotides with an annotated protein on the opposite strand ("prot. on opp. strand"), prediction on a scaffold for which no NCBI annotations were given ("unannot. scaff.") and all other predictions ("NA"). In all cases, most targets were highly covered by their MetaEuk prediction.

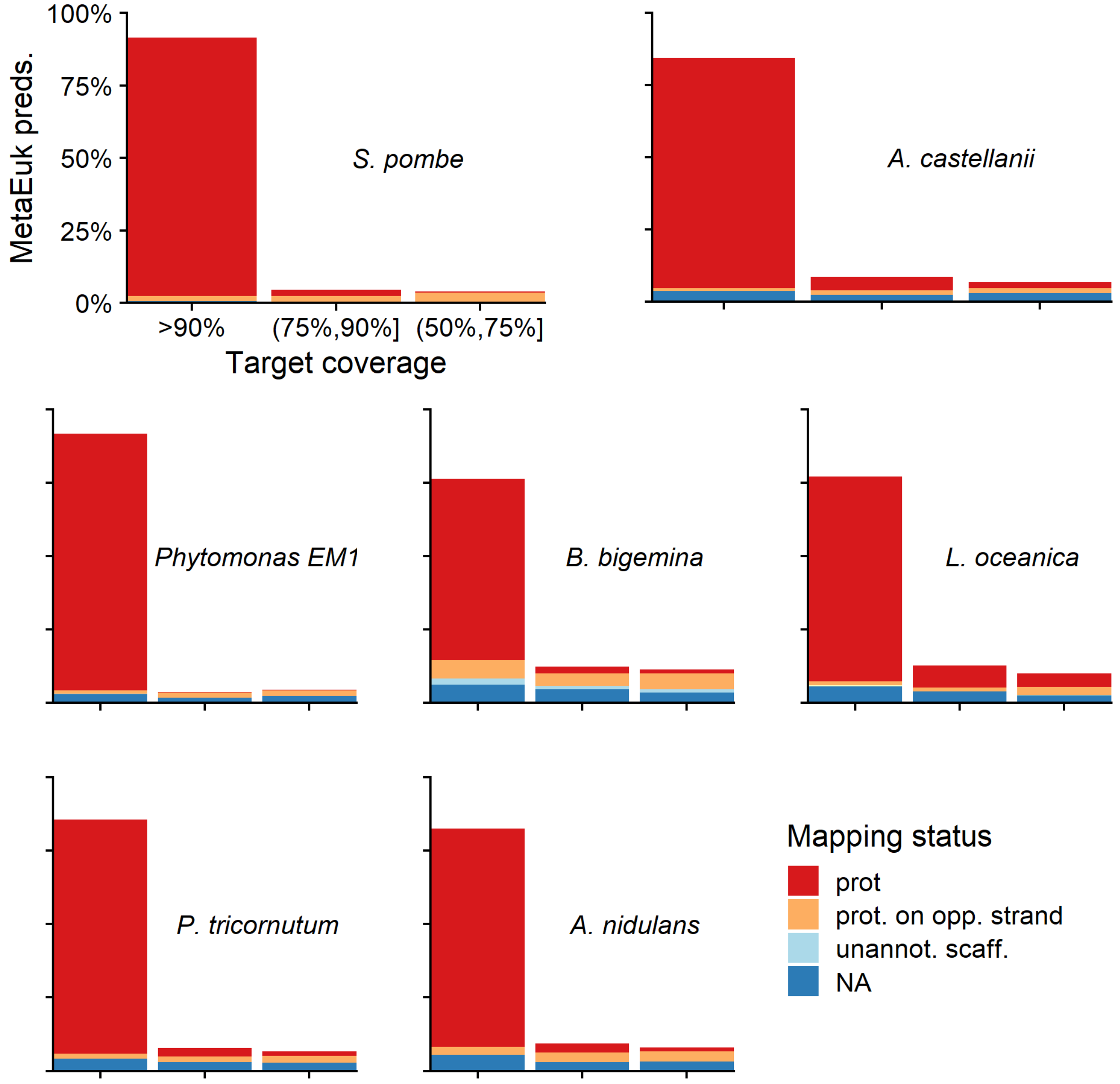

**Supplementary Figure 3 – MetaEuk E-values and bit-scores**

MetaEuk was run on a benchmark of seven eukaryotic unicellular organisms. The (A) E-values and (B) bit-scores computed between each predicted protein and its target by MetaEuk were compared to those computed by the Smith-Waterman algorithm. The Spearman rho values indicate high correlation for all benchmark organisms.

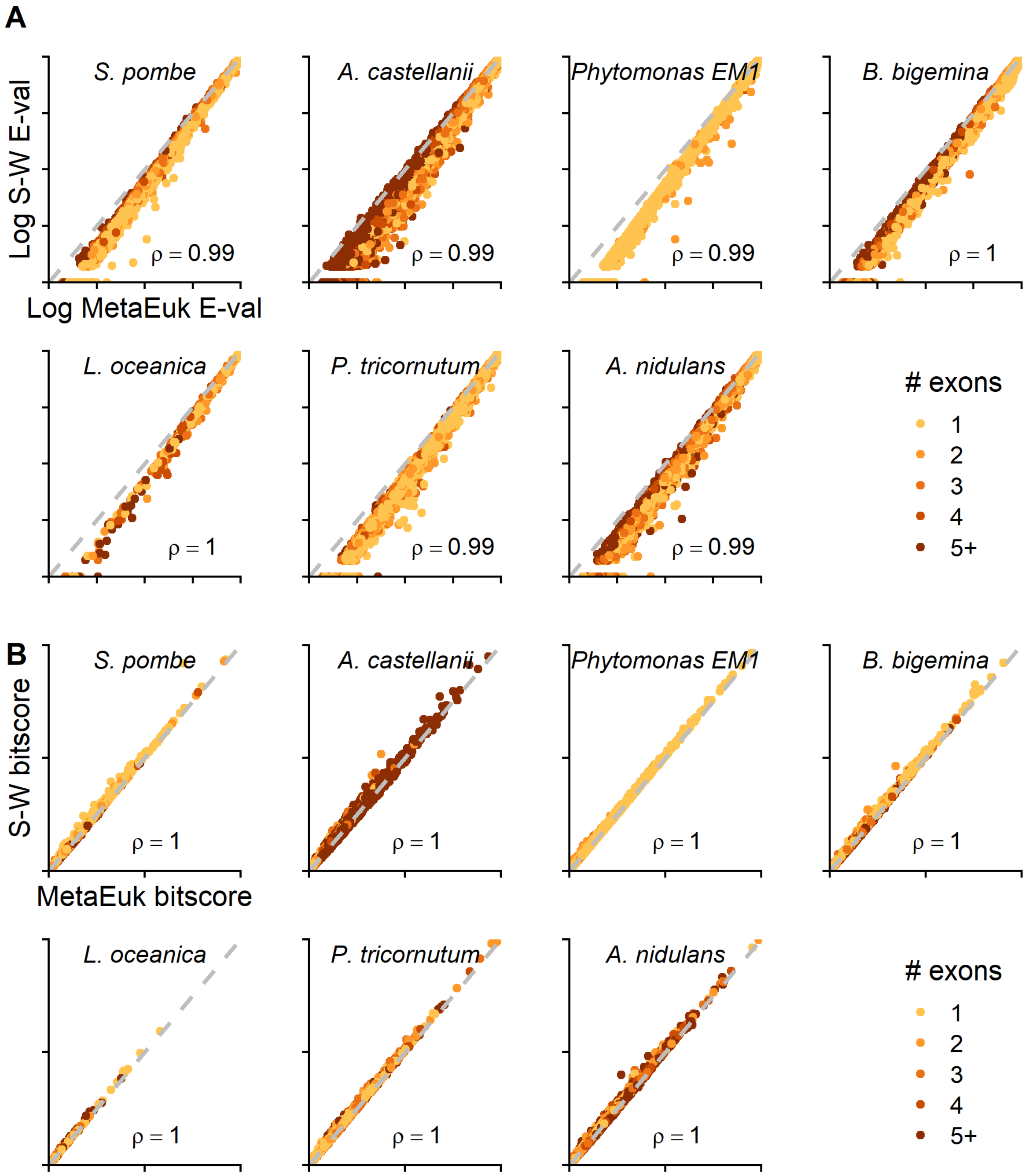

**Supplementary Figure 4 – MetaEuk evaluation on typical metagenomic contig lengths**

The annotated scaffolds of each of the organism in Table 1 were randomly divided into shorter contigs, following typical lengths of a metagenomics analysis (see Methods). Since each of the new contigs corresponds to locations on the original scaffolds, MetaEuk predictions on these contigs could be mapped to annotated proteins. (A) Conditions of increasing evolutionary divergence were simulated by excluding gene calls based on their sequence identity to their target. Sensitivity is the fraction of annotated proteins from the query genome to which a MetaEuk prediction was mapped. (B) Fraction of exons covered by MetaEuk (color saturation). The number of MetaEuk predictions is indicated on top of each bar. (C) In an annotation-dependent precision estimation MetaEuk predictions that mapped to an annotated protein were considered as “true” and the rest as “false”. (D) Fraction of annotated protein-coding genes that were split by MetaEuk into two (dark grey) or three (black) different predictions. (E) Comparison of the E-values computed by MetaEuk and by the Smith-Waterman algorithm for *A. castellani* proteins.

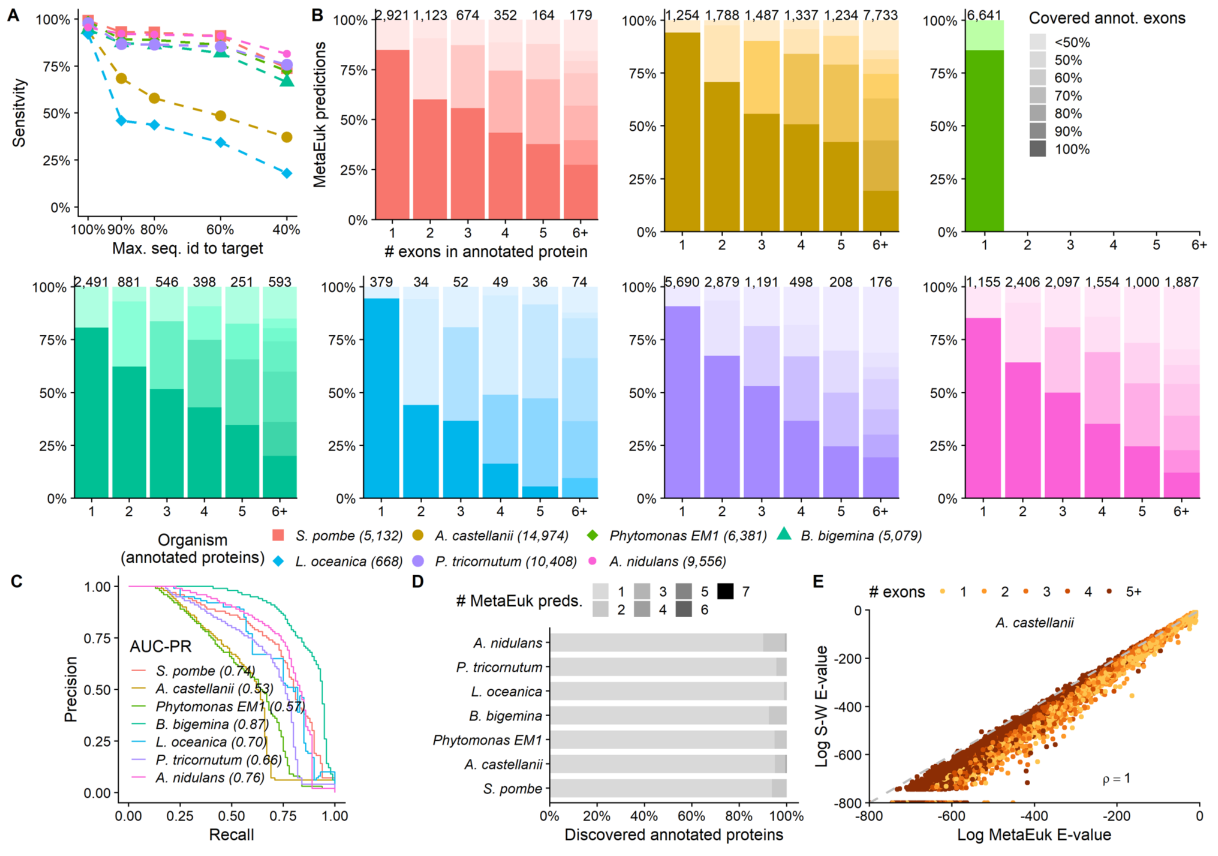

**Supplementary Figure 5 – Contribution of reference datasets to MetaEuk predictions**

Profiles computed based on clusters of MERC, MMETSP and Uniclust50 proteins served as the reference database for the MetaEuk run on the Tara Oceans contigs. MERC, MMETSP and Uniclust50 contributed 77.4%, 5.7% and 4.3% of the predictions, respectively. The rest of the predictions were based on mixed-dataset clusters.

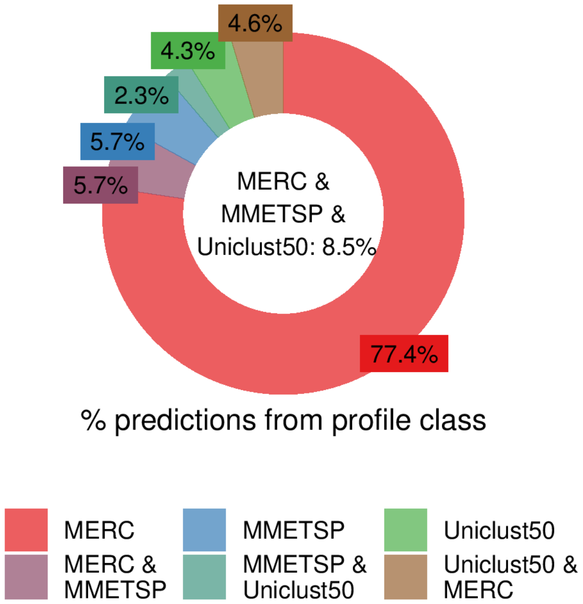
